# PKN is a sex- and species-specific fertilization factor in brown algae

**DOI:** 10.64898/2026.02.20.706836

**Authors:** Masakazu Hoshino, Meri Nehlsen, Rita A. Batista, Morgane Raphalen, Toshiyuki Wakimoto, Shinya Uwai, Kazuhiro Kogame, Vikram Alva, Susana Coelho

## Abstract

Fertilization, the union of male and female gametes, is central to sexual reproduction, yet the molecular mechanisms that ensure partner recognition and enforce species specificity remain elusive. Here we identify PKN, a previously uncharacterized transmembrane protein expressed exclusively in female gametes of brown algae, as an essential determinant of fertilization. Loss of PKN abolishes fertilization without affecting earlier mating steps, including gamete attraction, placing its function as a key mediator of male-female recognition. PKN contains extracellular β-propeller and mucin-like domains that are enriched in predicted glycosylation sites and rapidly evolving, consistent with a role in species-specific cell-cell recognition. Notably, PKN enforces reproductive isolation within the genus *Scytosiphon* by preventing interspecific fertilization. Together, these findings uncover a female-encoded recognition mechanism in brown algae and reveal protein-glycan interfaces as a conserved strategy for enforcing sex- and species-specific fertilization across Eukaryotes.

## Introduction

In sexually reproducing organisms, genetic information is transmitted across generations through the fusion of male and female gametes in a process known as fertilization. This event restores diploidy and initiates zygotic development. Despite its central role, fertilization is among the least understood fundamental biological processes, even in well-studied animal model systems, and the molecular mechanisms underlying many of its key steps remain elusive (*1*).

Fertilization generally follows a conserved sequence of events: chemotactic attraction between male and female gametes (or between cells of different mating types), gamete recognition, and ultimately, membrane fusion. The first step, attraction, is often mediated by chemical signals such as sex pheromones, released by female gametes, which guide male gametes, and ensure efficient fertilization (*2*, *3*). The second step, gamete recognition, involves highly specific molecular interactions between male and female gametes mediated by dedicated gamete recognition proteins (*4*, *5*). In many organisms, these recognition proteins are glycoproteins or lectins located on gamete surfaces, highlighting the broadly conserved importance of glycans, together with their protein backbones, as key determinants of recognition during fertilization (*4*, *6*). The final step, gamete fusion, requires the precise merging of the plasma membranes of the two gametes, allowing the mixing of haploid genomes (*7*, *8*). This process depends on specialized fusogenic proteins, such as the canonical HAP2/GCS1, a widely conserved eukaryotic fusogen that likely originated in the last eukaryotic common ancestor (LECA), with some notable lineage-specific losses (*9*).

Beyond enabling fertilization itself, gamete interactions are also central to species specificity and pre-zygotic reproductive isolation, preventing hybridization between closely related species (*10*). Accordingly, the molecular mechanisms governing gamete attraction and recognition are expected to also play a key role in mediating species-specific fertilization. This topic has attracted increasing attention, as it underpins both successful reproduction and the maintenance of species boundaries (*11*). Recent studies have begun to identify molecules involved in species-specific gamete interactions, ranging from the initial chemoattraction of sperm to the precise binding events at sperm-egg interfaces (*5*, *11*, *12*). Together, these discoveries suggest that gamete recognition molecules serve dual functions: promoting compatibility within species while simultaneously preventing cross-species fertilization. However, despite these advances, the molecular mechanisms that confer such extreme specificity remain poorly understood (*1*). Therefore, understanding how subtle variations in recognition proteins translate into strict species barriers remains a major challenge in reproductive and evolutionary biology.

Brown algae offer a unique and powerful system for investigating the molecular basis of gamete recognition and fertilization. Having diverged from the lineages leading to land plants and animals over a billion years ago, they represent an independent evolutionary trajectory in which complex multicellularity and diverse reproductive strategies have evolved (*13–15*). Their reproductive modes span isogamy, anisogamy, and oogamy, with fertilization typically occurring externally through free spawning (*15*). This reproductive ecology, combined with the relative ease of observing gamete interactions under the microscope, makes brown algae particularly well suited for dissecting the sequential steps of fertilization (*6*, *16*). Male gametes are attracted to females by pheromones, and females of a given species release a “bouquet” of different molecules, several of which are shared across species (*2*, *17*). Consistent with this, early work showed that these cues are relatively non-specific, since male gametes are often attracted to heterospecific female pheromones (*18*, *19*). This lack of specificity highlights the importance of subsequent steps, particularly gamete recognition and membrane fusion, in ensuring species specificity and reproductive fidelity (*20*). Several studies have shown that pre-incubation of gametes with lectins or specific sugars can strongly modulate fertilization efficiency (*24–26*). These observations led to the proposal that sex- and species-specific recognition may involve an interaction between a male gamete lectin and a glycoprotein on the female gamete membrane (*24–26*). Nevertheless, the nature of the factors mediating these interactions has remained elusive.

Here, we report the discovery of a novel transmembrane protein, which we name PICKINESS-ASSOCIATED PROTEIN (PKN). PKN is specifically expressed in female gametes and is required for male-female gamete recognition and, consequently, for successful fertilization. Structural analyses indicate that PKN contains multiple extracellular domains and numerous predicted glycosylation sites, consistent with a role in mediating cell-cell recognition. Using reverse genetics, functional assays and comparative analyses, we demonstrate that recognition mechanisms involving PKN are conserved across diverse brown algal lineages, highlighting the evolutionary significance of this protein. Importantly, we show that PKN confers species specificity to gamete interactions within the genus *Scytosiphon*, preventing interspecific fertilization and thereby ensuring reproductive fidelity. Together, these findings identify PKN as a key mediator of both sex- and species-specific gamete recognition and expand the repertoire of known eukaryotic fertilization proteins. More broadly, our results highlight sugar-protein interactions as a conserved and general mechanism for mediating gamete recognition and enforcing species-specific fertilization across domains of eukaryotic life.

## RESULTS

### A single locus controls conspecific fertilization preference in female gametes

We used the brown alga *Scytosiphon* (**Figure 1A**) for this study due to its capacity to produce large numbers of male and female gametes under laboratory conditions, as well as the availability of established genetic and genomic tools (*27–29*). *Scytosiphon* exhibits a heteromorphic alternation of generations, with a diploid multicellular sporophyte alternating with morphologically distinct haploid male and female multicellular gametophytes (*30*), **Figure 1A**). We carefully followed the mating ritual of these organisms: male gametophytes release numerous small, biflagellate gametes that actively swim, while female gametophytes produce slightly larger, similarly biflagellate motile gametes that quickly settle on the substrate. Fertilization occurs in three main steps: (i) once settled, female gametes secrete sex pheromones that chemoattract male gametes, which actively swim towards females; (ii) male gametes use their anterior flagella to scan the surface of female gametes until flagellum-membrane attachment is established; (iii) the cell bodies of the male and female gametes come into proximity and fusion occurs (**Figure 1B**).

**Figure 1.**
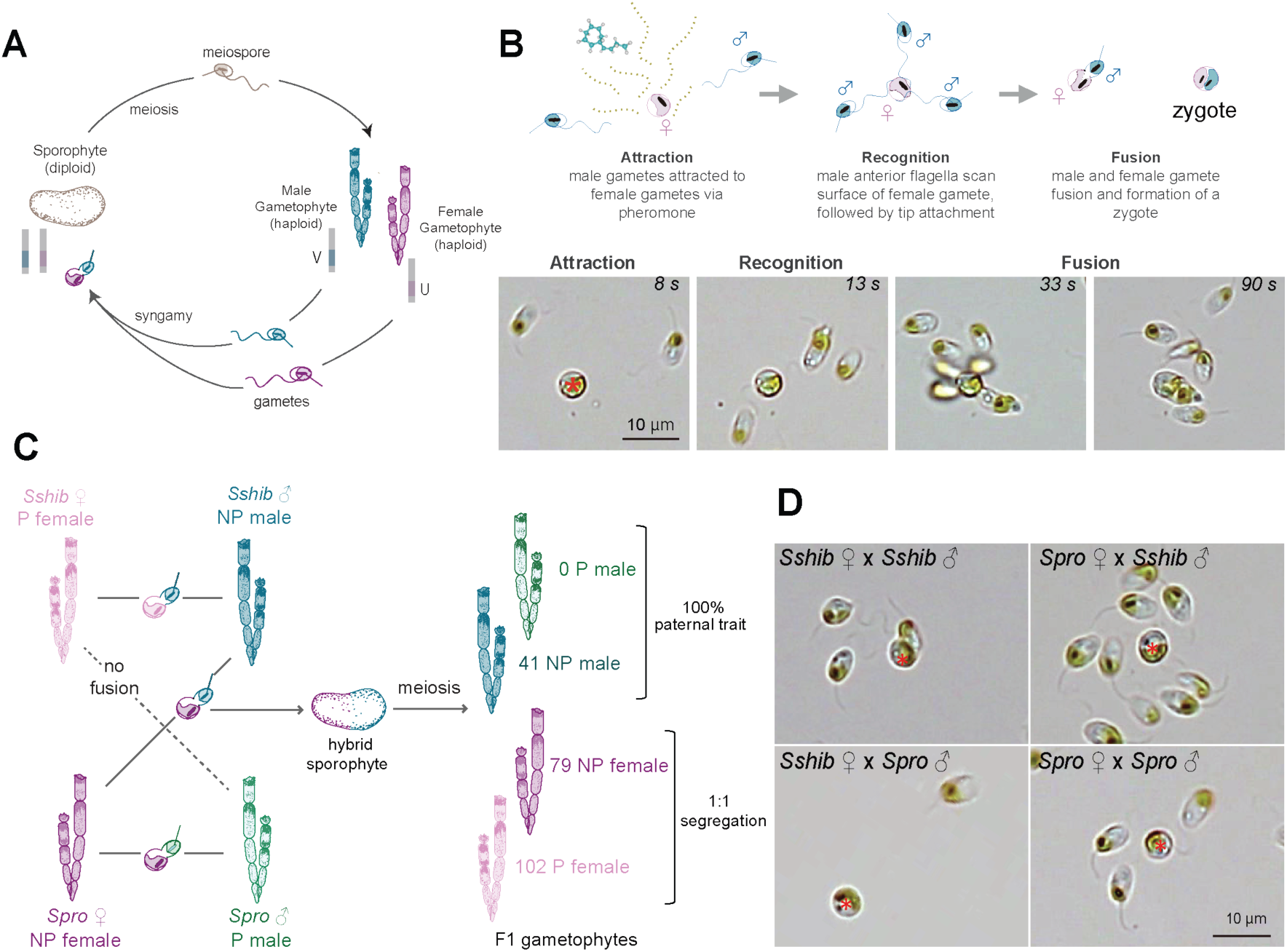
Life cycle and gamete interactions in the brown algae *Scytosiphon.* **(A**) Schematic diagram of the life cycle of *Scytosiphon*. Sex is determined at meiosis, depending on whether meiospores inherit a female (U) or male (V) chromosome. The sexual reproduction is nearly isogamous: female gametes are slightly larger than male gametes. (**B**) Gamete interaction during fertilization in *Scytosiphon*: illustration (upper) and time-lapse images (lower; female gametes are marked with red asterisks). The first step is the attraction of male gametes toward female gametes via pheromone production by the latter. Once male gametes reach the vicinity of a female gamete, they scan the female membrane and attach to its surface using their anterior flagella. The final step is contact between the cell membranes, followed by membrane fusion (**Video 1, 3**). (**C**) Schematic diagram of incomplete gamete incompatibility between *S. shibazakiorum* (*Sshib*) and *S. promiscuus* (*Spro*), and the phenotypes of F1 hybrid gametophytes. P: picky, NP: non picky. (**D**) Gamete incompatibility observed between *Sshib* female and *Spro* male. In the cross between *Sshib* female and *Spro* male, male gametes fail to attach their anterior flagella to the surface of female gametes. In contrast, in the other crosses, male gametes attach their anterior flagella to female gametes, resulting in multiple male gametes adhering to a single female via their flagella. Female gametes are indicated by red asterisks.

In order to investigate the molecular basis underlying species-specific fertilization, we focused on two *Scytosiphon* species, *S. promiscuus* (*Spro*) and *S. shibazakiorum* (*Sshib*), that exhibit asymmetric hybridization (*31*)(**Figure 1C**) Female gametes of *Spro* display a ‘non-picky’ (NP) phenotype, as they can be fertilized by male gametes of either species (**Videos 1, 2**). In contrast, female gametes of *Sshib* can be fertilized only by conspecific male gametes (**Videos 3, 4**), displaying a ‘picky’ (P) phenotype. This pickiness is associated with the inability of the anterior flagella of *Spro* male gametes to anchor to the plasma membrane of *Sshib* female gametes (**Figure 1D**; (*31*)).

To identify the locus involved in asymmetric hybridization, we produced a hybrid sporophyte resulting from the fertilization between *Spro* females and *Sshib* males, which at maturity produced haploid meiospores by meiosis (**Figure 1C**). Although these meiospores have a low survival rate (*31*), some successfully developed into gametophytes capable of producing gametes that could be backcrossed to the parental species. We therefore took advantage of one of these hybrid sporophytes to generate a segregating population of 578 individual gametophytes (**Table S1**).

We observed a 1:1 sex ratio of these F_1_ gametophytes (306 female and 271 male, **Table S1**). In total, 41 F_1_ males and 181 F_1_ females were also phenotyped for their ‘pickiness’ by backcrossing with both parental species (**Table S1**). Remarkably, all F_1_ males showed the same phenotype as their father (non-picky; i.e. their gametes fertilized female gametes of both species). In contrast, 1:1 segregation (*p* = 0.087; *G* test) of ‘picky’ and ‘non-picky’ phenotypes was observed in the F_1_ females (**Figure 1C; Table S1**). Specifically, 102 F_1_ females produced gametes in which the anchoring of the male anterior flagella (and subsequent fertilization) occurred only with *Sshib* males, but not with *Spro* males (P phenotype; **Video 5**). The remaining 79 F_1_ females produced gametes in which anchoring occurred with *Spro* males (NP phenotype; **Video 6**). Therefore, we conclude that the ‘pickiness’ of female gametes is determined by a single autosomal locus, which operates specifically in the female background.

### Identification of the locus responsible for female ‘pickiness’

In order to identify the autosomal locus involved in the female pickiness, we used a mapping-by sequencing-approach (see Methods). Using whole genome sequencing (**Table S2**) of 137 female F1 gametophytes (77 P and 60 NP), alongside with their parents, we identified a 253 kb region in chromosome 25 (position 1,965,159–2,218,259 bp) that was fully associated with the ‘pickiness’ phenotype of females (**Figure 2A-C**). PCR-genotyping of the region in 75 additional F1 females (**Table S3**) allowed to narrow down the candidate region to around 126 kbp (position 1,965,159–2,090,584 bp), where 13 genes are annotated (**Figure 2C**; **Table S4).**

**Figure 2.**
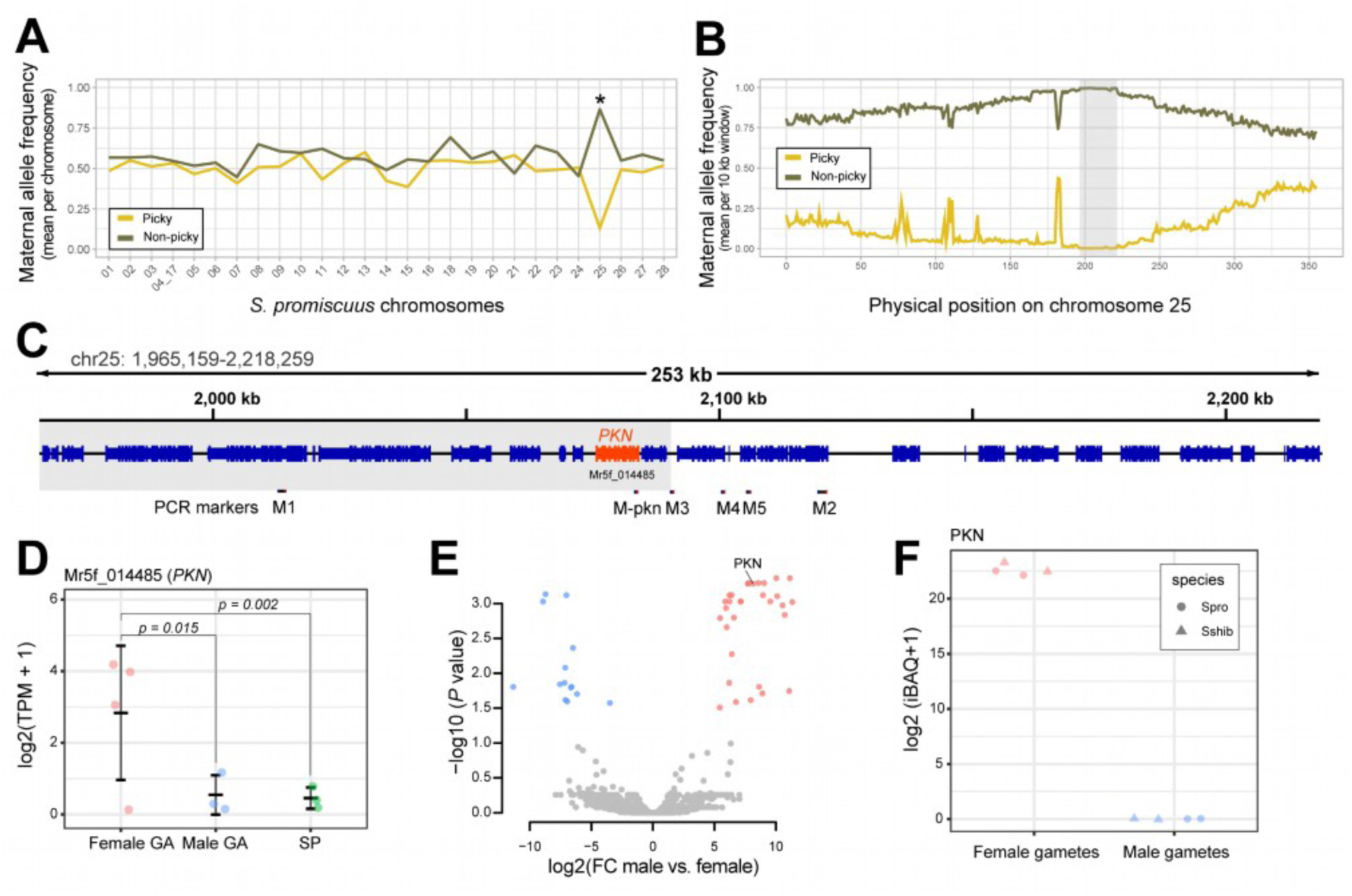
Identification of PKN in brown algae. (**A**) Average maternal allele frequency per chromosome among picky F1 females and non-picky F1 females. (**B**) Average maternal allele frequency on chromosome 25 (mean per 10 kb window) among picky F1 females and non-picky F1 females. The region associated with the phenotype (1.96–2.21 Mb) is highlighted in grey. (**C**) Annotated genes in the phenotype-associated region of chromosome 25 identified by WGS analysis. *PKN* is highlighted in orange, and other genes are in blue (exons as thick, tall bars; introns as thinner lines). Below the gene schematic, the positions of the six primers used for PCR-genotyping of individuals not included in the WGS analysis are indicated. The gray shaded area marks the genomic region linked to the phenotype, further refined by PCR-genotyping, containing 13 genes. (**D**) Gene expression level (log2 (TPM+1)) of *PKN* during the life cycle of *Spro*. GA: gametophyte, SP: sporophyte. Benjamin-Hochberg (BH) corrected *p*-values (< 0.05) are shown. (**E**) Volcano plot of differentially expressed proteins between female and male gametes. Male-biased proteins are shown in blue, and female-biased proteins in red (BH-corrected *p* < 0.05, |log₂ fold change| > 1). (**F**) log2 (iBAQ +1) values of PKN proteins in female and male gametes of *Spro* and *Sshib*. iBAQ: intensity-based absolute quantification.

To further refine the candidate gene(s), we examined the expression patterns of these 13 genes during the life cycle of *Scytosiphon*, reasoning that the gene involved in female pickiness would be preferentially expressed in female mature gametophytes, where gametes are produced. Transcriptomic analysis using RNA-sequencing (RNA-seq) across three life cycle stages (female and male gametophytes, as well as sporophytes) revealed that only one gene in the interval (gene Mr5f_014485) was markedly upregulated in female gametophytes (**Figure 2D**, **Figure S1, Table S4**). Finally, we used a proteomics approach to compare male and female gametes and identify sex-specific proteins (see Methods). Among the 18,787 proteins annotated in the genome, a total of 2,061 proteins were detected in three or more samples (**Table S5**). Differential proteomic analysis indicated that 33 were highly expressed in female gametes and 18 in male gametes (**Figure 2E**; **Table S5**). Among the 13 candidate genes identified through mapping-by-sequencing, only Mr5f_014485 was detected in this differential proteomic approach, and the protein was female-specific **(Figure 2E, F**).

Together, these results provide strong evidence that Mr5f_014485 is involved in the female-gamete selectivity (‘pickiness’) phenotype and therefore required for the species-recognition mechanisms in female gametes. We named the protein encoded by Mr5f_014485 PICKINESS-ASSOCIATED PROTEIN (PKN).

### Structure and evolution of the brown algal fertilization protein PKN

*Scytosiphon PKN* is predicted to encode a type I membrane protein (**Figure 3A**). This protein begins with an N-terminal signal peptide, followed by a 7-bladed β-propeller, a mucin-like intrinsically disordered region, a single transmembrane helix, and a short cytoplasmic tail enriched in acidic residues (Asp/Glu). An AlphaFold3 model of PKN reveals that the β-propeller exhibits a marked structural asymmetry: the distal (outer) face contains long, surface-exposed loop insertions, whereas the membrane-proximal (inner) face carries much shorter loops (**Figure 3B**). This architectural polarity suggests that the two sides may mediate distinct molecular interactions. Electrostatic surface calculations based on the AlphaFold3 model indicate that the outer face of the β-propeller carries a strongly negative potential, with a few small positively charged patches, whereas the inner face is only weakly negative by comparison (**Figure 3B**). Such a polarized electrostatic landscape is consistent with potential interactions with divalent cations (e.g., Ca²⁺) and/or positively charged protein partners, a feature commonly observed in propeller-mediated recognition modules. Furthermore, prediction of glycosylation sites suggested two putative N-glycosylated sites on the inner-face (**Figure S2**). The central mucin-like region is rich in Ser/Thr/Pro residues, typical of heavily O-glycosylated mucin domains. Consistent with this, abundant O-glycosylation sites were predicted in this region (**Figure S2A**), likely forming an extended, brush-like glycan coat. This sugar-rich region may function as a spacer that projects the β-propeller away from the membrane, a modulator of ligand accessibility, or a protective shield against proteolysis. Finally, the acidic cytoplasmic tail may mediate electrostatic contacts that contribute to PKN regulation or signaling.

**Figure 3.**
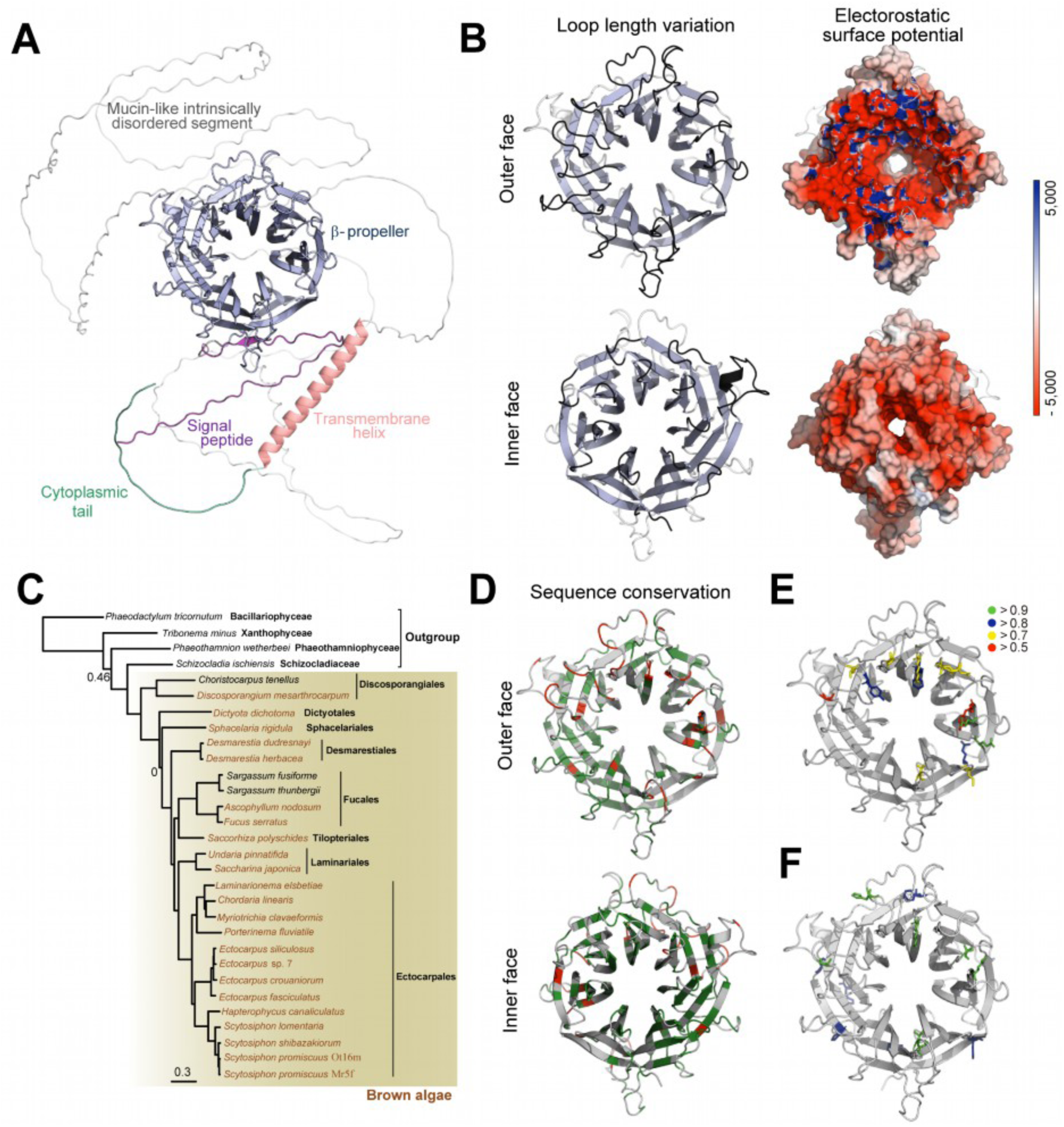
PKN structure and evolution. **(A)** AlphaFold3-predicted structure of PKN in Spro. (**B**) Characteristics of β-propeller of Spro PKN. Asymmetry of loop (highlighted in black) length and electrostatic surface potential (negative potential in red, and positive in blue) between outer and inner faces. (**C**) Species tree of brown algae and their close outgroups analyzed in this study, inferred by Orthofinder. Species in which PKN orthologues were identified are shown in brown. Superscript letters next to species names indicate the following: i, partial PKN orthologue detected; f, full-length PKN orthologue recovered but split across multiple contigs; d, multiple PKN orthologues were detected but they were partial. (**D**) Rate4Site fast-evolving residues in red and conserved residues in green. (**E**) Amino acid residues inferred to be under positive selection by PAML under the M8 model in codeml are shown in a line representation. Colors indicate posterior probabilities based on the BEB analysis: green for residues with probabilities ≥ 0.9, blue for ≥0.8, yellow for ≥0.7, and red for ≥0.5. (**F**) Amino acid residues that differ between *Spro* and *Sshib* PKN β-propeller. Residues on the distal face are in green and others in blue.

To infer the evolutionary origin of *PKN*, we examined genomes spanning brown algae and its close outgroups (Schizocladiaceae, Xanthophyceae, Phaeothamniophyceae, and Bacillariophyceae). Full-length *PKN* orthologues were identified in 18 brown algal species, including early-diverging groups such as Discospongiales and Dictyotales (**Figure 3C; Table S6)**. In several brown algal species, only partial *PKN*-like sequences were detected or no obvious orthologue was recovered (**Figure 3C; Table S6**), most likely reflecting incomplete genome assemblies, although lineage-specific gene loss cannot be entirely excluded. By contrast, no *PKN* orthologues were detected in any of the outgroup taxa. Consistently, BLASTP searches against the NCBI non-redundant (nr) database failed to detect *PKN* orthologues outside brown algae, suggesting that PKN is specific to this group of organisms. Within the genus *Scytosiphon*, PKN proteins share high pairwise sequence identity (86∼93%), whereas identity between *Spro* PKN and *Ectocarpus* PKN drops to ∼46%. Sequence identity is further reduced to ∼26% when *Spro* PKN is compared with PKN from the deeply branching species *Discosporangium mesarthrocarpum* (**Table S6**). Yet, despite this broad range of sequence divergence, all orthologues retain the same core domain architecture, including the characteristic β-propeller asymmetry and the strongly negative electrostatic potential on the outer face. In contrast, the central mucin-like disordered region varies markedly in length and composition across species, suggesting lineage-specific expansion or contraction of this segment while preserving the conserved structural core.

As signatures of rapid evolution in fertilization-related proteins are widespread across eukaryotes (*12*), we investigated whether PKN showsa similar patterns. Because PKN is highly divergent except for its β-propeller region, which is likely to mediate molecular interactions (*32*), we focused our analysis on this region. Distal (outer) loops of the β-propeller vary in length across the lineages (**Figure S3**). A Rate4Site analysis across all PKN orthologues further revealed that fast-evolving residues cluster on exposed loops of the distal surface, whereas the β-propeller core remains strongly conserved (**Figure 3D**). We also assessed positive selection by comparing rates of non-synonymous (*d_N_*) and synonymous (*d_S_*) substitutions across 17 brown algal species using CODEML implemented in PAML (*33*) (see methods). A likelihood ratio test (LRT) favored model M8, which allows sites with *ω* (*d_N_/d_S_*)) > 1 over the null model M7. Model M8 estimated that 6.1% of sites are under positive selection with *ω* = 1.74, and the Bayes Empirical Bayes (BEB) analysis identified 15 such sites with posterior probabilities greater than 0.5, including two with strong support (> 0.95: **Table S7**). All sites inferred to be under positive selection localize to the distal surface of the β-propeller (**Figure 3E**). In addition, the β-propeller domains of *Spro* and *Sshib* differ by 14 amino acid substitutions, seven of which map to the distal surface loops (**Figure 3F**), including two sites inferred as positively selected. Taken together, distal loop residues are among the most variable across PKN orthologues, suggesting that divergence in these exposed regions may broadly contribute to species-specific gamete recognition in brown algae.

### *PKN* is required for successful fertilization

To investigate the role of *PKN* during fertilization, we generated knockout (KO) *pkn* lines for female and male gametophytes of *Spro* and *Sshib*, using a CRISPR-Cas12 method (*29*). Three and six independent mutant lines were produced for female and male, respectively (**Figure 4A, Table S8**). *pkn* mutants of a *Spro* female (SproF-*pkn*2) and two *Spro* males (SproM-*pkn*1, SproM-*pkn*2) had an in-frame 6-bp deletion (i.e., two amino acid deletion) at the same position, which potentially disrupts proper folding of the 7-bladed β-propeller region (**Figure S4).** Mutations in other CRISPR lines led to a considerably disrupted predicted protein and a premature stop codon (**Figure 4A**).

**Figure 4.**
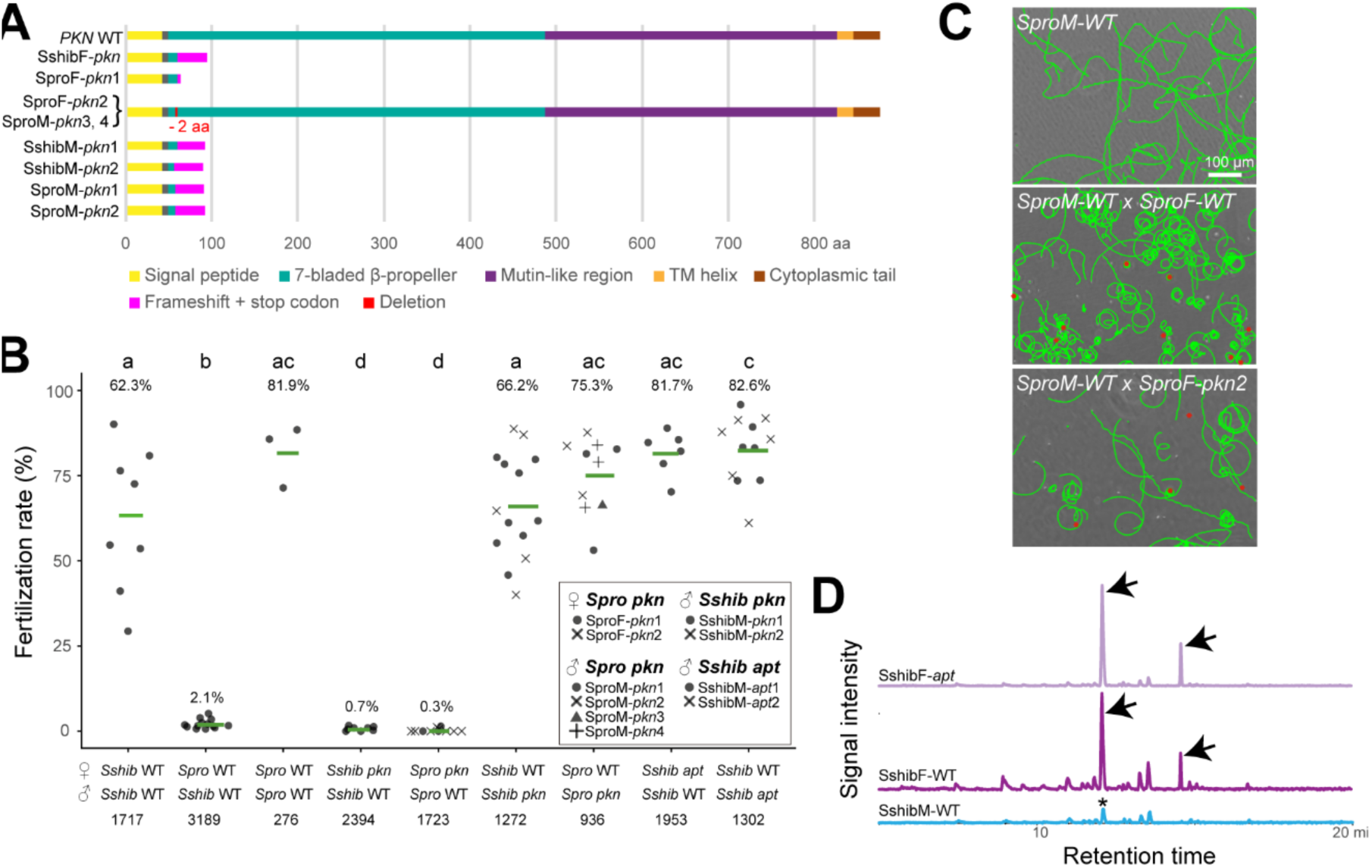
*PKN* mutagenesis. (**A**) Scheme of the wild-type (WT) PKN protein and the predicted proteins of each of the CRISPR-generated *pkn* mutants. (**B**) Fertilization rate of female gametes. Each point in the graph represents the fertilization success of an independent cross. The green horizontal lines and numbers above the plot show the mean fertilization rate. When multiple mutant strains were used in the crosses, different shapes were used to distinguish them. The different letters above the plots represent significant differences in mean fertilization rate (BH-corrected *p* < 0.05; Wilcoxon test: **Table S9**). Numbers below the plot represent number of scored gametes. (**C**) A 5 seconds long 2D trajectory of male gametes (green). WT male gametes only (upper), WT male gametes added to WT female gametes (middle), and WT male gametes added to *pkn* mutant female gametes (lower). Female gametes are indicated by red point. (**D**) Representative GC-MS analyses of the volatile compounds released from gametes: extracted ion chromatograms (MIC) = 79 *m*/*z*. The compound predicted as sex pheromone (ectocarpene, C_11_H_16_) was detected as two peaks (arrows) in WT and *pkn* mutant female.

In both *Spro* and *Sshib*, crosses between all female *pkn* mutants and wild-type males failed to produce zygotes **(Videos 7, 8**). In strike contrast, crosses between male *pkn* mutants and wild-type females yielded zygotes at levels comparable to control crosses (**Figure 4B**, **Table S9**, **Videos 9, 10**). Note that all mutant lines have an additional mutation at the *APT* locus that allows for selection CRISPR-modified lines (see methods; (*23*, *34*); nevertheless, the *apt* mutation has no effect on fertilization rates (**Figure 4B**, see also (*23*)). Together, these results indicate that loss of functional *PKN* causes infertility only when the mutation is present in a female background.

To determine which step of the mating pathway was disrupted in the mutants, we monitored the successive stages of fertilization in real time under a light microscope. Male gametes exhibit chemotactic behavior in response to the sex pheromone produced by female gametes (*23*). Upon detecting a decline in the pheromone concentration gradient, the posterior flagellum bends unilaterally, inducing a directional reorientation toward the source of pheromone emission (*35*). This results in a distinctive narrow circular trajectory of the gametes (**Figure 4C, middle)** that contrasts to the more linear trajectory when male gametes are alone (**Figure 4C, top)**. WT male gametes also show such narrow circular trajectory when they are crossed with female *pkn* mutants (**Figure 4C, bottom**), suggesting that the female *pkn* individuals secrets sex pheromone. Gas chromatography analysis (see Methods) confirmed that the *Sshib pkn* female mutant (SshibF-*pkn*) produces pheromone, thereby retaining the ability to attract male gametes (**Figure 4D**). These findings collectively suggest that the lack of fertilization is not attributable to impaired male-female attraction via the pheromone signal. Instead, our observations show that the anterior flagella of male gametes failed to attach to female *pkn* mutant gametes. Therefore, PKN is crucial for the recognition and physical interaction between the male flagella and the female gamete membrane, a prerequisite for subsequent approximation of the male and female cell bodies and gamete fusion (**Video 3**). Strikingly, this phenotype mirrors that of a ‘picky’ female *Sshib* when crossed with a male *Spro*, underscoring the critical role of PKN in mediating both male-female recognition and species-specific gamete interactions.

### PKN function is conserved in other brown algae

To test whether the function of PKN is conserved in other brown algae, we examined its role in *Ectocarpus* sp. 7, which also exhibits near-isogamous fertilization (**Figure 5B, Video 11**) but diverged from *Scytosiphon* >10 million years ago (*13*, *21*, *27*, *36*). Using CRISPR-Cas9 and 12, we generated two independent female gametophyte mutant lines, EcF-*pkn1* and EcF-*pkn2* (**Figure 5A**, **Table S11**). EcF-*pkn1* carries a 2 bp deletion in the first exon, producing a premature stop codon and a truncated 92-amino-acid protein, whereas EcF-*pkn2* harbors a 343 bp deletion removing part of the open reading frame, including the start codon. Both lines, like previously reported for *Sshib* and *Spro* CRISPR mutants, include a mutation in the *APT* locus for selection of CRISPR-modified lines Crosses of wild-type male gametes with mutant female gametes failed to produce zygotes in both lines (**Figure 5C**). Male gametes appeared to be attracted to mutant females, but their anterior flagella failed to attach to the female surface, recapitulating the phenotype observed in *Scytosiphon* (**Figure 5 D, Videos 12, 13).** These results demonstrate that PKN is required for male-female gamete recognition in two evolutionarily distinct brown algal species, highlighting its conserved role in sexual reproduction.

**Figure 5.**
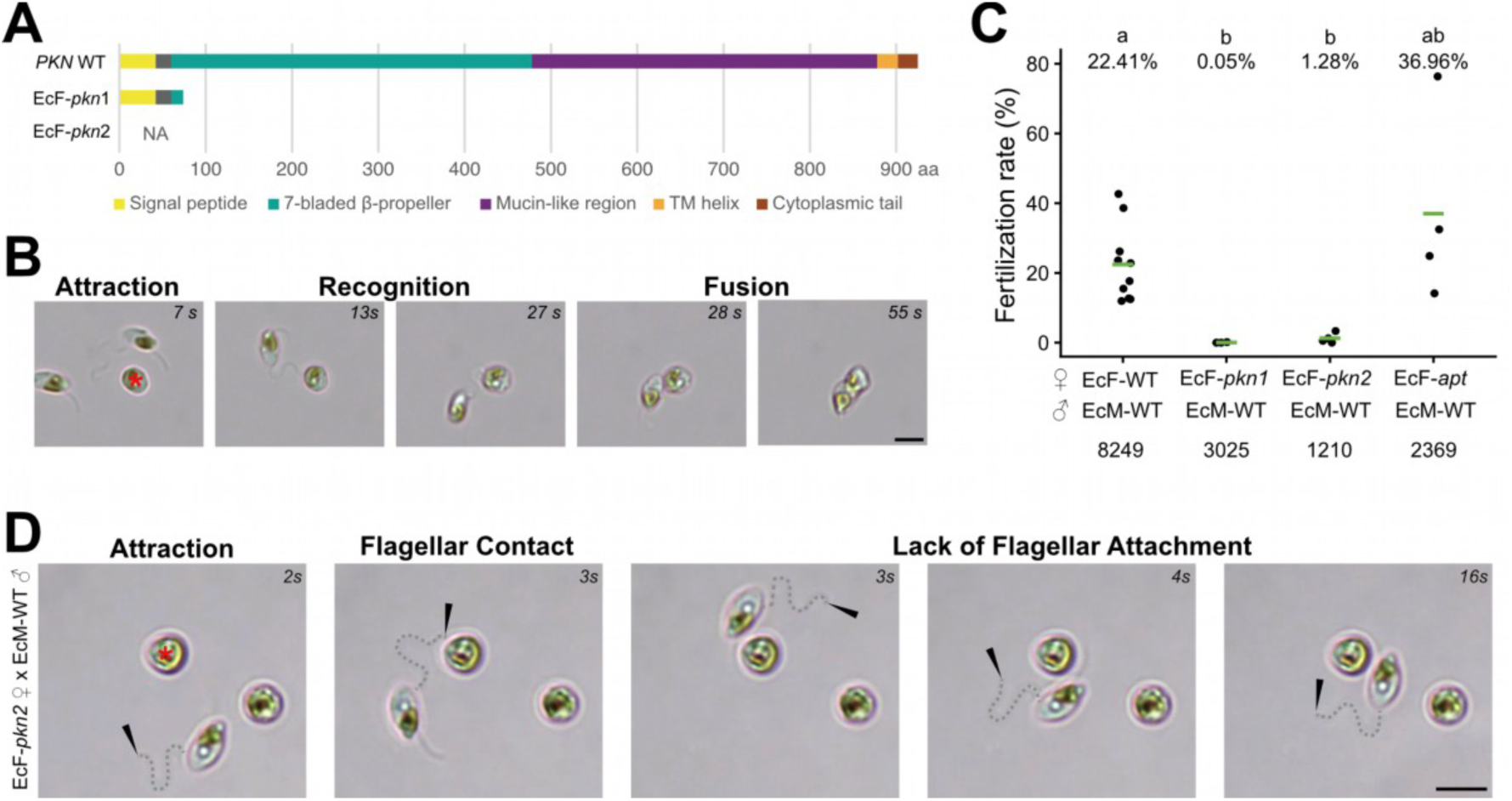
Role of PKN in *Ectocarpus* fertilization. (**A**) Scheme of *Ectocarpus* wild-type (WT) and *pkn* mutant predicted proteins (**Table S11**). (**B**) Gamete interaction during fertilization in *Ectocarpus.* The red asterisk denotes the female gamete. The male circles the female gamete, attracted by production of the sex pheromone, before anchoring to the surface of the female with the tip of its longer anterior flagellum. Upon attachment, male gametes display a rapid beating of their anterior flagella (activation) and after the gametes fuse. (**C**) Fertilization rate of WT and *pkn* mutant female gametes. Note that the *pkn* mutant females *EcF-pkn1* and *EcF-pkn2* also carry the same mutation in the APT gene as *EcF-apt* mutants. Each point represents the fertilization rate of an independent cross. The green horizontal lines and numbers above the plot show the mean fertilization rate. Numbers below the plot indicate the number of female gametes screened. Letters above the plot represent significant differences in the mean fertilization rate (BH-corrected p < 0.05; Wilcoxon test: **Table S12**). Numbers below the plot represent number of scored gametes. (**D**) Impaired fertilization in female *pkn* mutants. When crossed with *pkn* mutant females, male gametes displayed a chemotatic response, but while the tip of the anterior flagellum (as indicated by the dashed grey line and the black arrows) contacted the surface of the female gamete, it did not attach. Instead, males continued to circle the female gametes without activating and fusing (**Videos 12, 13**).

## Discussion

Our experiments identify PKN as a key genetic determinant of gamete recognition in brown algae. PKN is expressed specifically in female gametes, and the protein localizes to the female gamete surface. Functional analyses show that loss of PKN in female gametes abolishes fertilization, due to failure of the male anterior flagellum to anchor to the female gamete surface.

Remarkably, PKN has been co-opted to mediate species-specific compatibility in *Scytosiphon*: the anchoring defect mirrors the gametic incompatibility observed in interspecific crosses between *Sshib* females and *Spro* males. Genetic mapping confirms that PKN is fully linked to this gametic incompatibility, indicating that the same protein underlies both male-female recognition and species-level reproductive isolation in this genus.

Interestingly, our phenotypic assays in *Ectocarpus* show that PKN plays a comparable role in female-male recognition and that this process is similarly mediated by male flagellum anchoring in the female gamete membrane. This observation, together with previous reports showing that male flagellar anchoring is highly species-specific and fails in interspecific *Ectocarpus* crosses (2) suggests that PKN has been repeatedly co-opted to enforce gamete recognition and species specificity across brown algae.

PKN encodes an extracellular β-propeller domain followed by a mucin-like region, both likely mediating interactions with ligands on the anterior flagellum of male gametes. Both domains contain multiple predicted glycosylation sites, consistent with earlier studies suggesting that female gamete recognition proteins in brown algae are glycoproteins (*24*, *25*). Glycoprotein-mediated recognition is a recurrent feature of fertilization systems across eukaryotes (*4*, *37*), underscoring the broad evolutionary importance of protein-glycan interactions in reproductive processes. For instance in fish, the egg membrane glycoprotein Bouncer is required for sperm-egg binding and restricts fertilization to conspecific sperm in certain species combinations (*11*), with differences in amino acid composition and N-glycosylation patterns proposed to contribute to species specificity (*38*).

In PKN, predicted N-glycosylation sites within the β-propeller are conserved between *Spro* and *Sshib*, suggesting that glycosylation alone does not explain gamete incompatibility between these species. By contrast, the outer-face loops of the β-propeller show rapid evolution and substantial variation in length and sequence across lineages, implicating them as determinants of species-specific receptor function. Consistent with this, multiple amino acid substitutions were observed in these loop regions between *Spro* and *Sshib*, raising the possibility that they underlie gamete incompatibility between the two species. While the functional role of individual residues cannot yet be tested experimentally, due to the lack of transgenic tools in brown algae, the evolutionary pattern strongly implicates the β-propeller surface as a key specificity-determining interface.

PKN represents the first gamete recognition protein identified in brown algae. Its orthologues are broadly distributed across this lineage and show signatures of positive selection, consistent with the rapid evolution observed for gamete recognition proteins in other eukaryotes (*12*, *39*). Remarkably, PKN is conserved in brown algal species spanning diverse reproductive modes, from isogamy to oogamy, highlighting a pervasive role in fertilization despite substantial variation in gamete morphology and mating systems. Functional conservation over at least ∼10 million years, as inferred from our experiments with the *Ectocarpus* orthologue, strongly supports the idea that PKN-mediated gamete recognition is a longstanding and essential feature of brown algal reproduction. At the same time, PKN shows no detectable homology to known gamete recognition proteins in any other eukaryotes and appears restricted to brown algae, identifying it as a lineage-specific innovation.

Taken together, PKN exemplifies a novel female-encoded gamete recognition protein that both mediates male-female interactions and reinforces species boundaries in a major eukaryotic lineage. Its discovery provides a unique opportunity to understand how fertilization systems evolve in multicellular lineages that reproduce via broadcast spawning, where the interplay of molecular specificity and environmental promiscuity shapes reproductive success.

## Methods

### Production and phenotyping of F_1_ hybrids

F_1_ hybrid gametophytes between *Spro* female (strain Mr5f) and *Sshib* male (strain Os10m) were produced following (*31*)(**Table S1**). The sex of the F_1_ strains weas verified using PCR-based sex markers (*40*) and test crosses. To phenotype F_1_ strains, they were crossed with the parental species, and the occurrence of anchoring between the male anterior flagellum and the female plasma membrane was assessed. F_1_ females were classified as having a picky (P) phenotype if anchoring was established with *Sshib* males, but not with *Spro* males. Conversely, females were classified as non-picky (NP) phenotype if anchoring was observed with *Spro* males. Similarly, F_1_ males were classified as P if anchoring occurred with *Spro* females but not with *Sshib* females, whereas they were classified as NP if anchoring was observed with *Sshib* females. Crossing experiments were performed as in (*31*), and culture strains used are listed in **Table S1**.

### DNA extraction and sequencing for the segregation family

Although the parental strains and 137 F_1_ female gametophytes had already been sequenced in a previous study (*41*), we conducted additional genome sequencing on the parental strains and five F1 female gametophytes to increase the number of samples and sequencing depth (**Table S2**). Prior the genome DNA extraction, the gametophytes were cultivated with antibiotics (Penicillin G 50 mg/L, Ampicillin 50 mg/L, Chloramphenicol 5 mg/L, and Kanamycin 0.5g/L) for one week to remove bacterial contamination. The gametophytes were frozen in liquid nitrogen and crushed to powder using TissueLyser II (QIAGEN, Venlo, Netherlands) and genomic DNA was extracted using OmniPrep^TM^ for Plant (G-BIOSCIENCES, St. Louis, USA), following the manufacture’s instruction. Libraries were prepared following (*42*) and sequenced on an illumina NextSeq2000 with 150-bp paired-end reads.

### Genome assembly and genome annotation

To assemble the genomes of the mother (*Spro*, Mr5f) and father strain (*Sshib*, Os10m), raw reads were trimmed and filtered with Trimmomatic v0.39 (*43*) and subsequently assembled with Platanus Assembler v1.2.4 (*44*). Contigs originated from contaminated bacteria were identified using Blobtools v1.1.1 (*45*) and manually removed. Remaining contigs were then scaffolded by RagTag v2.0.1 (*46*) using a chromosome-level genome assembly of *S. promiscuus* (*14*). Because both maternal and paternal genome assemblies had many gaps, some of them were closed by performing PCR and Sanger sequencing (primers are listed in **Table S6**). The scaffolded genome was then annotated by Funannotate v1.8.17 pipeline (*47*). To annotate the genomes, publicly available RNA-seq data of *Spro* was used for the maternal genome, while RNA-seq was newly performed for the paternal genome (**Table S10**). RNA was extracted from mature female gametophytes of *Sshib* (strain Os1f), which cultivated at 10°C short day condition (lignt:dark, 8 h:16 h). Thalli were frozen in liquid nitrogen and crushed to powder using Bead crusher μT-01 (TAITEC, Saitama, Japan) and total RNA was extracted using ISOSPIN Plant RNA kit (Nippon Gene Co. Ltd., Tokyo, Japan), and the extract was purified using RNA Clean & Concentrator-5 (Zymo Research, Carifornia, USA). An RNA library was then prepared using NEBNext Ultra II Directional RNA Library Prep with Sample Purification Bead (New England Biolabs, Massachusetts, USA), and was sequenced on an illumina NovaSeq 6000 platform with 150-bp paired-end reads.

### Identification of the genomic region linked to the ‘pickiness’ of female gametes

To identify the genomic region linked to the pickiness of female gametes, we analyzed maternal allele frequency of each phenotype groups. Previously generated genome-seq data of the two parental and 138 F1 female gametophytes (77 P and 61 NP female) were retrieved from GenBank (**Tables S2)**. Raw reads of each individual were trimmed and filtered with Trimmomatic v0.39 and then mapped onto the maternal genome using BWA v0.7.17 with the BWA-MEM (*48*).The resulting *sam* files were converted to *bam* files and sorted by coordinates using Samtools v1.6 (*49*). To call SNPs, the sorted *bam* files were analyzed with GATK v4.6.2.0 pipeline (*50*). SNPs with depth below 3 and allele frequency below 0.3 were filtered out, and the maternal or paternal origin of each SNP was identified in each individual using VCFtools v0.1.16 (*51*). The average maternal allele frequency was calculated at every 10 kb window and visualized for each phenotype group with R software v4.5.0.

To narrow down the genomic region identified above, six PCR markers were designed within the region (**Figure 2C**), and genotyping was performed on 75 phenotyped F1 female individuals that were not used for genome sequencing (**Table S3**).

### Transcriptome and proteome analysis

To investigate the expression levels of genes encoded in the genomic region linked to the pickiness of female gametes, transcriptome and proteome analyses were performed. Publicly available *Spro* RNA-seq data (female gametophyte, male gametophytes, and sporophytes: **Table S10**) was analyzed with the nf-core/rnaseq pipeline v.3.14.0 (*52*), using the genome of the mother strain as the reference, and STAR-Salmon alignment and quantification strategy (*53*, *54*). The resulted transcripts per million (TPM) values for each gene were visualized with R software. Deferential expression (DE) between samples was assessed using DESeq2 v1.42 (*55*).

For proteome analysis, female and male gametes of both *Spro* and *Sshib* (two replicates for each: total eight samples) were mixed with loading buffer and individually loaded onto a NuPAGE 12% Bis-Tris Gel (Thermo Fisher Scientific, Massachusetts, US) for a short SDS based gel electrophoresis and stained with colloidal Coomassie using the ReadyBlue Protein Gel Stain (Merck Millipore, Massachusetts, US). Gel sections containing proteins were excised, and cut into smaller parts. For in-gel digestion of proteins, gel pieces were destained by washing three times with 5 mM ammonium bicarbonate (ABC) in acetonitrile (ACN) (1:1, v/v) for 20min. After a dehydration step with 100% ACN for 10min, disulfide bonds were reduced with 10 mM dithiothreitol (DTT) in 20 mM ABC for 45 min at 56°C, and thiol groups of cysteine residues were prevented from reoxidation by carbamidomethylation with 55 mM iodoacetamide (IAA) in 20mM ABC for 45 min in the dark. Gel pieces were then washed two times with 5 mM ABC in ACN (1:1, v/v) for 20min and dehydrated with 100% ACN for 15min. After evaporation of the liquid in a vacuum centrifuge for 10 min, gel pieces were soaked in a solution of 12.5 ng/µl sequencing grade trypsin (Promega) in 20 mM ABC, pH 8.0 for 10 min at room temperature (RT), and then covered with 20 mM ABC. After in-gel digestion of proteins at 37°C overnight, peptides were extracted in three consecutive steps with different extraction buffers for 30 min: first 3% (v/v) trifluoroacetic acid (TFA) in 30% (v/v) ACN was added, followed by 0.5% (v/v) formic acid (FA) in 80% (v/v) ACN, and finally by 100% ACN. ACN was evaporated from pooled supernatants by vacuum centrifugation. In the course of the digestion protocol all incubation steps were carried out under shaking.

Peptides were desalted with C_18_ StageTips (*56*) and analyzed on an EASY-nLC 1200 UHPLC coupled to a Q Exactive HF mass spectrometer (both Thermo Fisher Scientific) as described previously (*57*) with little modification: peptides were separated on the analytical column using a 60 min segmented gradient of 10-33-50% of HPLC solvent B (80% acetonitrile in 0.1% formic acid) at a flow rate of 200 nl/min.

In the mass spectrometer, MS and MS/MS spectra were generated at resolution 60k. Full MS target value and maximum IT were set to 3×10^6^ and 25 ms, respectively. In each scan cycle, the 7 most intense precursor ions were picked up. The MS/MS target value was set to 10^5^ charges with a maximum IT of 220 ms.

The MS data from all replicates were processed together using MaxQuant software suite v.2.2.0.0 (*58*). Database search was performed using the Andromeda search engine (*59*), which is integrated in MaxQuant. MS/MS spectra were searched against a target-decoy database consisting of 18787 protein entries, which were annotated in the genome of *Spro* strain Mr5f, and 245 commonly observed contaminants. In database search, full specificity was required for trypsin. Up to two missed cleavages were allowed. Carbamidomethylation of cysteine was set as fixed modification, whereas oxidation of methionine and acetylation of protein N-terminus were set as variable modifications. Initial mass tolerance was set to 4.5 parts per million (ppm) for precursor ions and 0.5 dalton (Da) for fragment ions. Peptide, protein and modification site identifications were reported at a false discovery rate (FDR) of 0.01, estimated by the target/decoy approach (*60*).

The abundances of the detected proteins were quantified using iBAQ (intensity-based absolute quantification). Proteins detected in fewer than three samples were excluded from the downstream statistical analyses. iBAQ values were log2-transformed after adding a pseudocount of 1 to avoid undefined values. Subsequently, missing values (i.e., zeros) were imputed with the minimum non-zero value observed across the dataset. DE analysis between female and male gametes was conducted using the limma package v3.64.1(*61*) in R software. A linear model was fitted with both biological condition (i.e., sex) and batch (reflecting measurement day) as factors to account for potential batch effect.

In both transcriptome and proteome DE analyses, genes/proteins with an adjusted *p*-value (BH corrected) < 0.05 and absolute log2FC > 1 were considered significantly differentially expressed.

### Evolutionary analysis and structural models of *PKN*

*PKN* orthologues were identified using OrthoFinder v3.0.1b1 (*62*) and BLASTP (*63*). For OrthoFinder, the proteome of the mother strain (*Spro*), which includes PKN, was searched against proteomes of 23 brown algal species and four ochrophytes outgroups (Schizocladiaceae, Xanthophyceae, Phaeothamniophyceae, and Bacillariophyceae), with default settings. These proteome datasets comprised both publicly available sequences and newly annotated proteomes generated from existing genome assemblies (**Table S6**: see below for details of the annotation procedure). For BLASTP, *Spro* the PKN protein was queried against the NCBI non-redundant (nr) database and the above brown algal dataset. Candidate sequences were retained if they contained the characteristic signal peptide, seven-bladed β-propeller domain, mucin-like region, and C-terminal transmembrane helix. Structural models for all orthologues were generated using the AlphaFold3 webserver (*64*), and domain boundaries, loop architecture, and surface electrostatics were examined in PyMOL. Furthermore, for *Scytosiphon* PKN, O- and N-glycosylation sites were predicted using NetOGlyc v4.0 (*65*) and NetNGlyc v1.0 (*66*). Only sites with a NetOGlyc score of ≥ 0.99 and sites with full Jury agreement (9/9; NetNGlyc score) were considered as potential O- and N-glycosylation sites, respectively.

For phylogenetic analysis, β-propeller regions were extracted from the AlphaFold3 models and aligned using MAFFT (*67*) with the L-INS-i strategy (mafft--maxiterate 1000--localpair). Maximum-likelihood trees were inferred using IQ-TREE v3 (*68*) with automatic model selection (MFP), 1,000 ultrafast bootstrap replicates, and 1,000 SH-aLRT tests. WAG+F+I+G4 was selected as the best-fitting model according to BIC, and the obtained tree was visualized in iTOL (*69*). Site-specific evolutionary rates were calculated using Rate4Site (*70*), using the maximum-likelihood tree generated by IQ-TREE as the guide tree. Rate4Site was run in empirical Bayesian mode with 16 discrete rate categories (-ib -k 16), and the resulting scores were mapped onto the structural models in PyMOL, where negative scores correspond to conserved sites and positive scores reflect rapidly evolving residues. Divergent and conserved positions identified by Rate4Site were mapped onto the β-propeller structure in PyMOL by selecting residues with high (b > 1.5) or low (b < –0.5) relative rate scores. Positive selection was also assessed for 15 species, for which complete sequence of *PKN* β-propeller domain were available, using the CODEML program implemented in PAML v4.10.9 (*33*, *71*). CODEML was performed following the procedure described by (*72*), using the mutation–selection model of codon frequency (*73*) and two site models: M8, which allows for positively selected sites, and its null model M7, which does not (*74*). The M8 and M7 models were compared by the LRT, and positively selected sites were identified by BEB method (*75*).

Prior to the *PKN* orthologues identification, most of the publicly available genomes of brown algae and Schizocladiaceae were re-annotated. Genome assemblies downloaded from public databases were annotated using either the Funannotate v1.8.17 pipeline (*47*) with publicly available RNA-seq data (**Table S10**) or the GALBA v.1.0.11 pipeline (*76*) with proteomes from closely related taxa. Gene models of annotated *PKN* orthologues were then manually curated. For *Scytosiphon lomentaria*, because the previously published genome assembly (*28*) was highly fragmented, we reassembled the genome using previously published reads following the same procedure applied to the parental strains.

### CRISPR-mediated mutagenesis and phenotyping of *PKN* mutants

Using a CRISPR-Cas12 system, we knocked out the *PKN* gene in both male and female individuals of *Spro* and *Sshib*, following the protocol described in (*29*). We used a single selection gene encoding adenine phosphoribosyl transferase (*APT*), which generates a resistant phenotype when algae are cultured in a medium with 2-fluoroadenine. Thus, all *pkn* mutants isolated have a mutation at the *APT* locus. Mutations were confirmed by PCR and Sanger sequencing. Guide RNAs for *PKN* and *APT* were designed using CRISPOR web server (*77*). Sequences of the Guide RNAs and PCR primers are described in **Table S8**.

The fertilization rate of *pkn* mutants was quantified following the protocol described in (*78*), with WT and *apt* mutants examined as controls. The fertilization rate of each experimental group was compared by pairwise Wilcox rank sum test followed by the Benjamini-Hochberg correction method. Interaction between female and male gametes during fertilization was observed under either an inverted light microscope (CKX53: Olympus, Tokyo, Japan) or a compound light microscope (BX51: Olympus), and video recordings were made using a video camera: Floyo-200 (Wraymer Inc., Osaka, Japan) for the inverted light microscope, and BC1200 (Sato Shouji Inc., Kawasaki, Japan) for the compound light microscope.

Production of sex pheromone by *pkn* mutants were tested by observing the trajectory of male gametes and using a gas chromatography-mass spectrometry (GC-MS). It is known that male gametes of brown algae exhibit characteristic swimming trajectories in the presence of sex pheromones (*79*). Thus, the swimming trajectories of WT male gametes against *pkn* female mutant gametes were examined. Male gametes were added to the settled female gametes, and their movement was recorded with video camera Floyo-200 under the inverted light microscope. The trajectory of male gametes were visualized using TrackMate (*80*) implemented in Fiji (*81*).

For GC-MS, mature gametophytes of *Sshib* female *pkn* mutant (SshibF-*pkn*) were kept in 200 mL of sterilized seawater in a 300 mL flask at 10°C and gametes were released in the flask. Volatile secretions were trapped on Mono Trap RCC18 (GL science, Tokyo, Japan) using a closed-loop-stripping system. After looping for 16 h at 10°C, absorbed volatile compounds were eluted with 50 μL of CH_2_Cl_2_ and immediately analyzed by GCMS. [Zebron ZB-wax (phenomenex) 30 m × 0.25 μm; He as the carrier gas; program rate: 45-200°C at 5°C/min]. Compounds were identified by using the NIST MS library and similarity search program. Mature gametophytes of WT female and male of *Sshib* were also examined as positive and negative controls.

### CRISPR-mediated mutagenesis and phenotyping of *Ectocarpus pkn* mutants

As described for *Scytosiphon* we generated CRISPR mutants for the *PKN* gene alongside *APT* as a selection gene in female individuals of *Ectocarpus* sp.7. Sequences of the guide RNAs and PCR primers used are available in **Table S11**.

In order to quantify fertilization rates in *Ectocarpus*, five to 10 mature gametophytes were collected and dried, placed in a dry 35-mm petri dish, and incubated in the dark at 14 °C for 3-4 hours. The gametophytes were then flooded with a 500-700 µL drop of ice-cold natural sea water (NSW) and placed in front of a directional light source. Shortly after, swimming gametes could be pipetted from the side of the NSW drop that faced the light. To quantify the fertilization rate, flow channels of a chambered coverslip (µ-Slide VI, 80606: ibidi, Gräfelfing, Germany) were filled with 30 µL NSW before adding 3-7 µL of swimming female gametes, which were left to settle in the dark at 14 °C for 20 - 30 minutes. Unsettled females were then removed by washing the channel with NSW. To determine the number of settled female gametes the slide was imaged using an inverted light microscope (Observer Z1: ZEISS, Oberkochen, Germany) and a camera (Axiocam 702 mono: ZEISS, Oberkochen, Germany) and images were analyzed using FIJI (ImageJ2 version 2.16.0/1.54p). Then, 3-7 µL of swimming male gametes were pipetted from the light-facing side of the NSW drop, added to the channels and left at 14 °C overnight. Zygotes defined by the presence of two eyespots (*82*, *83*) were manually counted under an inverted light microscope, and the fertilization rate was estimated as the number of zygotes divided by the number of settled female gametes (*84*). The fertilization rates of *pkn* mutants were compared to both wild-type and *apt* mutants as a control by pairwise Wilcoxon rank sum test with a Benjamini-Hochberg correction for multiple testing.

To record the fertilization of the *pkn* mutants and wild-type controls, gametes were released and added to a chambered coverslip as described above for the quantification of the fertilization rate. Videos of the fertilization events were recorded using an inverted light microscope (Axiovert 5: ZEISS, Oberkochen, Germany) and a colour camera (DFK 37AUX250: The ImagingSource, Bremen, Germany).

## Acknowledgements

We thank Marek Kuka and Frank Chan for assistance in the library preparations and helpful discussions. We thank Ana Velic and Boris Maček from the proteomics facility at the University of Tübingen. We also thank Jun Ishigohoka, Josué Barrera-Redondo, and Sodai Lotharukpong, for discussion regarding data analyses. This work was supported by the Max Planck Gesellschaft, the ERC (grant n. 638240 to SMC), the Moore Foundation (GBMF11489 to SMC) and the Bettencourt-Schuller Foundation. MH was supported by JSPS KAKENHI Grant number 23K19386, JSPS Overseas Research Fellowships, and the Max Planck Gesellschaft.

## Supplemental Videos

**Video 1.** Intraspecies crosses in *Spro*. Male gametes were added to settled female gametes. Strain As8f x As6m.

**Video 2.** Interspecies crosses between *Spro* female and *Sshib* male: strain As8f x Os10m.

**Video 3.** Intraspecies crosses in *Sshib*: strain Os8f x Os10m

**Video 4.** Interspecies crosses between *Sshib* female and *Spro* male. At first, *Spro* male gametes (strain As6m) were added to settled *Sshib* female gametes (strain Os8f), and subsequently *Sshib* male gametes (strain Os10m) were added to make sure that the *Sshib* female gametes are fertile.

**Video 5.** Fertilization phenotype of F1 picky female. At first, *Spro* male gametes (strain Mr8m) were added to settled female gametes of F1 picky female (strain MO181), and subsequently *Sshib* male gametes (strain Os10m) were added.

**Video 6.** Fertilization phenotype of F1 non-picky female. *Spro* male gametes (strain Mr8m) were added to settled female gametes of F1 non-picky female (strain MO257).

**Video 7.** Fertilization phenotype in crosses of *Spro pkn* females (Spro♀-*apt/pkn*1) with wild-type *Spro* males (Spro♂-WT).

**Video 8.** Fertilization phenotype in crosses of *Sshib pkn* females (Sshib♀-*apt/pkn*1) with wild-type *Sshib* males (Sshib♂-WT).

**Video 9.** Fertilization phenotype in crosses of wild-type *Spro* females (Spro♀-WT1) with *Spro pkn* males (Spro♂-*apt/pkn*4).

**Video 10.** Fertilization phenotype in crosses of wild-type *Sshib* females (Sshib♀-WT) with *Sshib pkn* males (Sshib♂-*apt/pkn*2).

**Video 11.** Fertilization phenotype in crosses of *Ectocarpus* wild-type females (EcF-WT) with wild-type males (EcM-WT).

**Video 12.** Fertilization phenotype in crosses of *Ectocarpus pkn* females (EcF-*pkn1*) with wild-type males (EcM-WT).

**Video 13.** Fertilization phenotype in crosses of *Ectocarpus pkn* females (EcF-*pkn2*) with wild-type males (EcM-WT).

## Supplemental Figures

**Figure S1.**
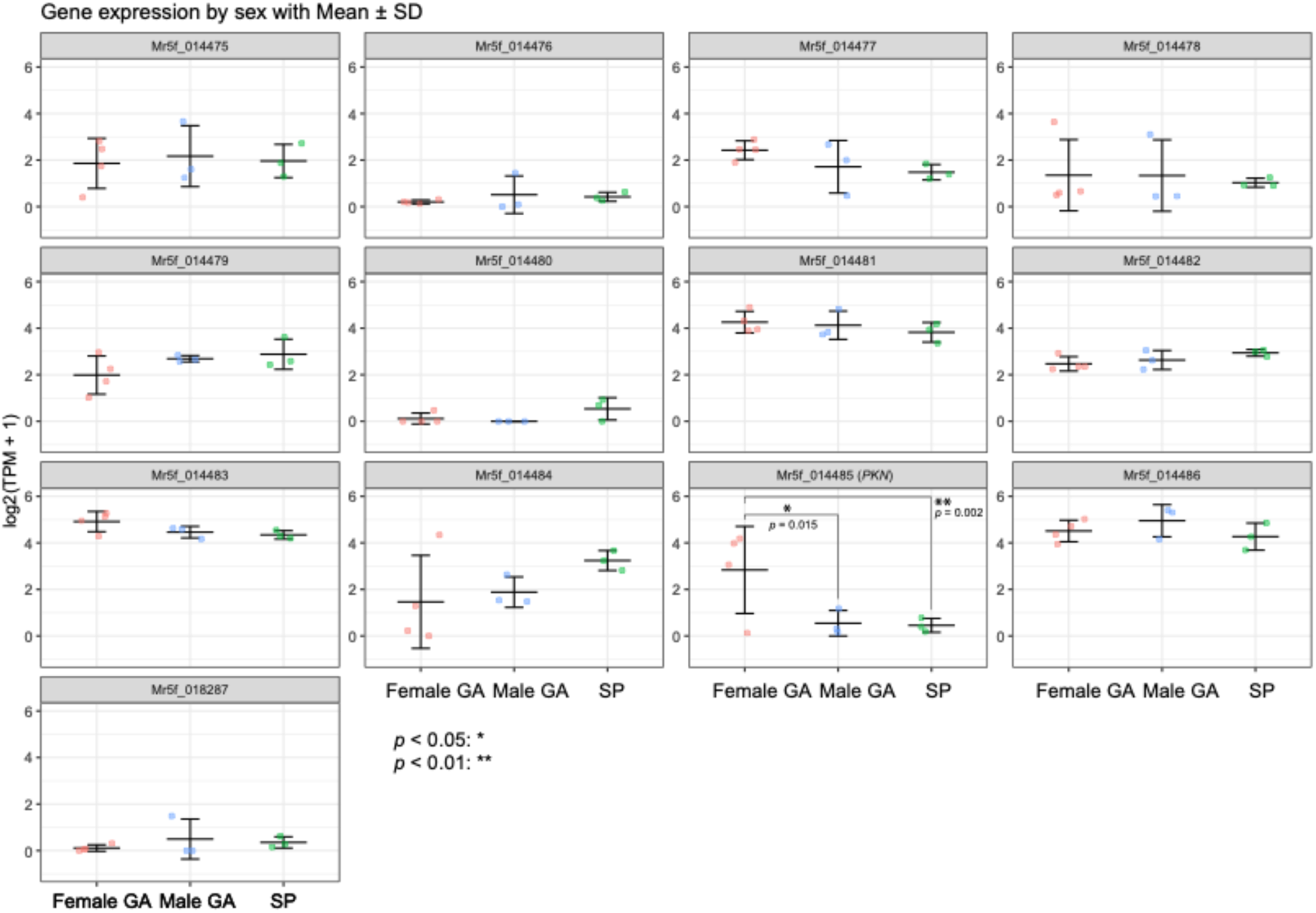
Gene expression level (log2 (TPM+1) of the 13 genes, which are annotated on the genomic region linked to the pickiness of female gametes, during the life cycle of *Spro*. Only *PKN* (Mr5f_014485) has female-biased expression. Exact *p*-values for each gene are listed in **Table S4**. GA: gametophyte, SP: sporophyte. Benjamin-Hochberg (BH) corrected *p*-values (< 0.05) are shown.

**Figure S2.**
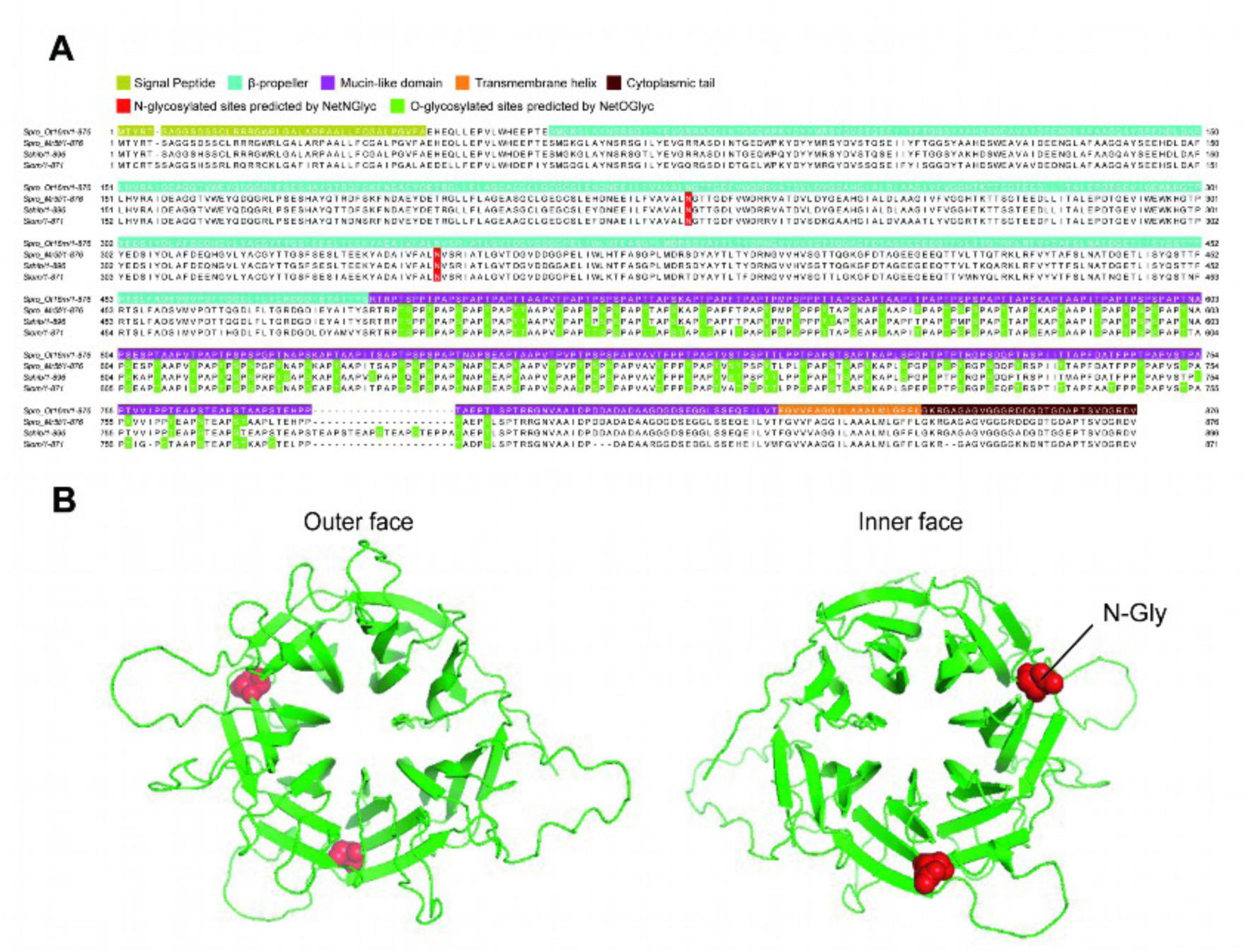
Predicted glycosylation sites on the *Scytosiphon* PKN. (A) Alignment of the four PKN sequences of *Scytosiphon*. From top to bottom: the sequence of *Spro* strain Ot16m with domain annotation, the sequence of *Spro* strain Mr5f, the sequence of *Sshib*, and the sequence of *S. lomentaria*. In the bottom three sequences, possible N-glycosylation sites are shown in red and O-glycosylation sites in green. (B) Two predicted N-glycosylation sites, represented as red spheres, on the β-propeller of *Spro* PKN (strain Mt5f).

**Figure S3.**
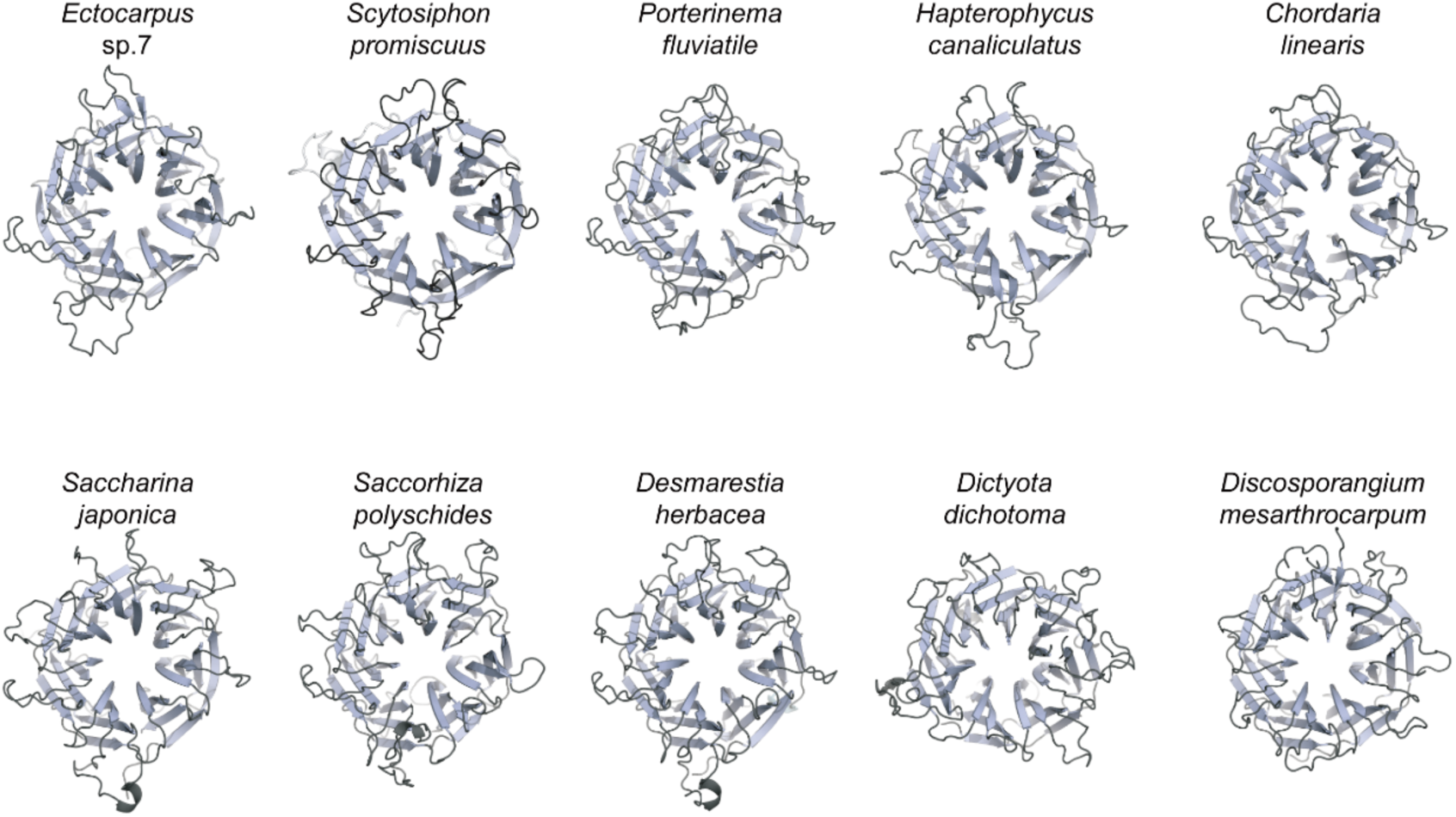
Variation in the loop length of the distal face of the β-propeller among various brown algae. The loop regions are highlighted in black.

**Figure S4.**
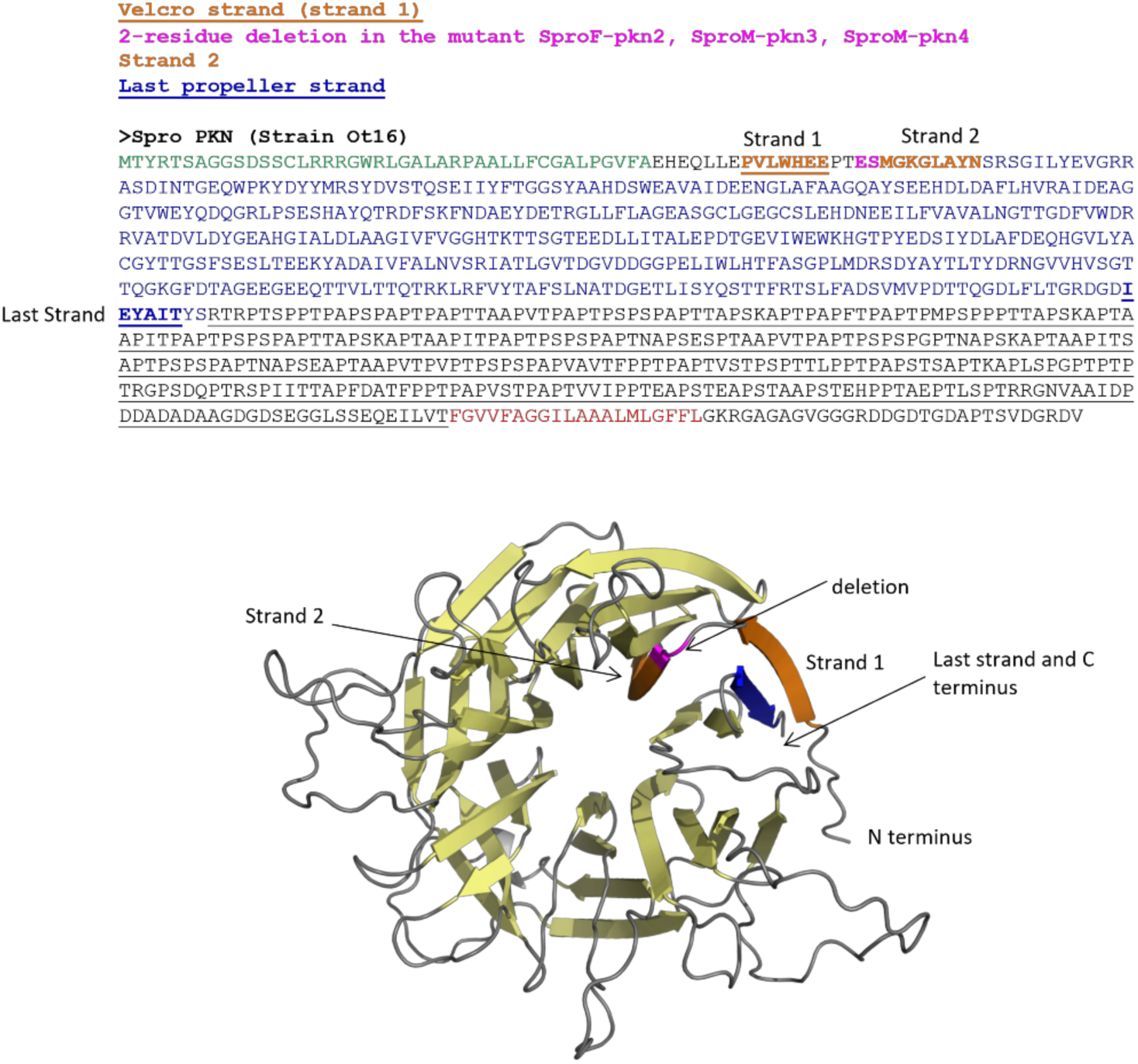
Position of the two amino acid deletion in the *Spro pkn* mutants: SproF-*pkn*2, SproM-*pkn*1, SproM-*pkn*2. In the β-propeller fold, the first β-strand interacts with the last one to form a “velcro” closure. The deleted residues are in the loop connecting the first and second beta-strands, and it is likely that the deletion disrupts the proper folding of the propeller by influencing the interaction between the first and last β-strands.

